# Erythroferrone Modulates Osteoblast-Osteoclast Crosstalk During Bone Remodeling

**DOI:** 10.64898/2026.05.05.722997

**Authors:** Pinanong Na-Phatthalung, Gabrielle van Caloen, Marina Planoutene, Emily Tai, Anisa Gumerova, Georgii Pevnev, Jay Cao, Ronit Witztum, Eva Ingber, Leon Kautz, Farhath Sultana, Funda Korkmaz, Maayan Levy, Tony Yuen, Mone Zaidi, Yelena Z. Ginzburg

## Abstract

Erythroferrone (ERFE) secretion inhibits hepcidin expression by sequestering several bone morphogenetic protein (BMP) family members to increase iron availability for erythropoiesis. Recent evidence demonstrates that ERFE is also expressed in osteoblasts and osteoclasts and *Erfe*^−/−^ mice display low–bone–mass arising from increased bone resorption despite a concomitant increase in bone formation. To mechanistically dissect how bone–derived ERFE exerts an osteoprotective effect, we first created *Erfe*^*fl/fl*^ mice, which were then crossed with *Col2.3*-Cre mice to generate osteoblast–selective *Erfe* mutants (or *Erfe*^*fl/fl*^;*Col2.3*-Cre mice). We now demonstrate that ERFE derived from osteoblasts is not responsible for the decreased BMD noted in *Erfe*^−/−^ mice, revealing enhanced BMD during anabolic stress in *Erfe*^*fl/fl*^;*Col2.3*-Cre mice. Consistently, in contrast to global ERFE loss, osteoblast–selective ERFE loss does not increase osteoclasts *in vivo*. Furthermore, our results demonstrate that ERFE loss in osteoblasts induces osteoclast *Erfe* expression in co-culture experiments *in vitro*. Finally, the osteoclastogenesis gene program is induced in co-culture with osteoblasts only when ERFE is lost in osteoclasts. Taken together, our finding provide strong evidence of osteoclast–derived ERFE as a central osteoprotective regulator of bone mass, its loss resulting in net bone loss in *Erfe*^−/−^ mice.

**BRIEF SUMMARY:** Loss of erythroferrone derived from osteoclasts enhances osteoclastogenesis resulting in accelerated bone loss.

**SIGNIFICANCE STATEMENT:** Canonical erythroferrone (ERFE) function includes hepcidin suppression through bone morphogenic proteins (BMPs) sequestration, establishing the rationale for ERFE-mediated regulation of bone homeostasis. We previously showed that global ERFE loss controls bone mass. Here, we report that osteoclast–derived ERFE is a major regulator of osteoclastogenesis. For this, we crossed *Erfe*^*fl/fl*^ with *Col2.3*-Cre mice to generate osteoblast–selective *Erfe* mutants, demonstrating that osteoblast–derived ERFE does not recapitulate bone loss found in global ERFE knockout mice. In contrast, bone mineral density is enhanced during anabolic stress in *Erfe*^*fl/fl*^;*Col2.3*-Cre mice. Finally, we document that osteoclast ERFE loss enhances osteoclastogenesis in co-culture with osteoblasts. Together, the data provide compelling evidence that osteoclast–derived ERFE modulates communication between osteoblasts and osteoclasts.

## INTRODUCTION

Erythroferrone (ERFE) was first described as an erythroblast-derived hormone regulating the crosstalk between erythropoiesis and systemic iron supply (1). Specifically, ERFE was identified as a central physiological erythroid regulator of hepcidin, the principal regulator of systemic iron metabolism (2-4). Hepcidin prevents iron egress from cells by binding to and blocking the iron exporter ferroportin (FPN1) (5-7). Hepcidin expression is induced by iron loading and inflammation and suppressed by hypoxia, iron deficiency, and expanded or ineffective erythropoiesis (6, 8, 9). Importantly, iron stores regulate hepcidin by several bone morphogenic proteins (BMPs) and ERFE regulates hepcidin expression by sequestering BMPs (10, 11), which results in dampened BMP/SMAD signaling (12).

We previously demonstrated that, because BMPs are crucially important for bone modeling and remodeling (13), and ERFE function involves BMP sequestration (10, 11), ERFE is also centrally involved in bone remodeling. Most notably, we discovered that osteoblasts and osteoclasts express *Erfe* mRNA, that *Erfe*^−/−^ mice exhibit low bone mineral density (BMD) with increased number of osteoclasts, and that global ERFE loss enhanced bone resorption (14). Specifically, *Erfe*^−/−^ osteoblasts exhibit enhanced *Sost* and *Rankl* expression and BMP–mediated signaling *ex vivo*, increasing osteoclastogenesis in *Erfe*^−/−^ mice. These findings are consistent with recent literature (15-19) demonstrating that loss of BMP signaling increases bone mass through direct osteoclast inhibition. Furthermore, we found that osteoblasts isolated from *Erfe*^−/−^ mice displayed enhanced mineralization and activation of the osteoblastogenesis gene program and postulated that enhanced osteoblast activity further enhanced osteoclast activity, resulting in net bone loss in *Erfe*^−/−^ mice (14).

As a metabolically active endocrine organ, bone constantly remodels in response to changing metabolic and mechanical needs (20). Osteoblasts arise from mesenchymal stem cells and their production, activity and survival is regulated by complex circuitry involving multiple hormones and cytokines, including the BMPs and Wnts (21). In contrast, osteoclasts are derived from hematopoietic stem cells, primarily under the control of M-CSF and RANKL (20). Given that BMPs are critical for bone development, modeling and remodeling (21) and that ERFE action involves BMP sequestration (10, 11, 22), we hypothesized that also ERFE coordinates bone homeostasis by titrating BMPs to regulate net bone mass —determined by a balance between osteoblastic bone formation and osteoclastic bone resorption — by coordinating the crosstalk between osteoblasts and osteoclasts.

On the basis of these findings, we proposed to dissect the mechanisms through which bone ERFE regulates bone remodeling. To accomplish this, we generated *Erfe*^*fl/fl*^ mice. Previously, using *Erfe*^*fl/fl*^;*Col2.3*-Cre mice, we reported that ERFE derived selectively from osteoblasts is a critical driver of systemic hepcidin suppression and impedes hemoglobin recovery during phlebotomy–induced stress erythropoiesis *via* a BMP4–dependent mechanism (23). This study revealed a critical role of osteoblast–derived ERFE in the regulatory crosstalk between erythropoiesis and iron metabolism. The purpose of the current effort is to fully characterize the effects of osteoblast–selective ERFE loss on bone homeostasis and dissect mechanisms thereof. We now show that ERFE derived from osteoblasts is not responsible for the decreased BMD noted in *Erfe*^−/−^ mice, revealing enhanced BMD during anabolic stress in *Erfe*^*fl/fl*^;*Col2.3*-Cre mice. Consistently, in contrast to global ERFE loss (14), osteoblast–selective ERFE loss does not increase osteoclasts *in vivo*. Furthermore, our results demonstrate that ERFE loss in osteoblasts induces osteoclast *Erfe* expression in co-culture experiments *in vitro*. Finally, the osteoclastogenesis gene program is induced in co-culture experiments only when ERFE is lost in osteoclasts. Taken together, our finding provide strong evidence of osteoclast–derived ERFE as a central osteoprotective regulator of bone mass, its loss resulting in net bone loss in *Erfe*^−/−^ mice.

## MATERIALS AND METHODS

### Mice

*Erfe* global knockout mice (*Erfe*^−/−^) were generous gifts from Tomas Ganz, UCLA (1). C57BL/6 (wild type) mice and osteoblast type Iα1 collagen 2.3-kb promoter-cre mice (*Col2.3*-Cre) were purchased from Jackson Laboratories. The floxed *Erfe* mouse line was produced by GEIiC (Genome Engineering & iPSC Center, Washington University in St Louis). Osteoblast-specific *Erfe* knockout mice (*Col2.3*-Cre;*Erfe*^fl/fl^) were generated by crossing *Col2.3*-Cre mice with floxed *Erfe* mice. The targeted *Erfe* allele and cre recombinase gene were identified by PCR using tail biopsy with specific primer pairs listed in Table S1. The Institutional Animal Care and Use Committee at Icahn School of Medicine at Mount Sinai approved all animal experimental procedures. All mice had access to food and water ad libitum under AAALAC guidelines.

### DEXA and Micro-CT scanning

Skeletal phenotyping was conducted on 6 weeks and 14 weeks old mice male unless otherwise indicated. Bone mineral density (BMD) of the whole body, femur, tibia, and lumbar vertebrae (L4-L6) was measured in sedated mice using dual energy X-Ray absorptiometry (DEXA) (Scintica instrumentation, Scintica Inc., TX, USA). To stimulate bone remodeling, 6 weeks old mice were injected with 50 µg/kg human parathyroid hormone (PTH) (#H-4835.0005, Bachem) or vehicle (0.9% sodium chloride) daily for 8 days and sacrificed 48 hours after the last injection. To stimulate bone loss, ovariectomy was performed in 20 weeks old female mice to induce estrogen depletion; ovariectomized mice were sacrificed 20 weeks after surgery. After mice were sacrificed, femur, tibia, and lumbar vertebrae (L4) were collected, cleaned, and fixed in 10% neutral buffered formalin (#5735, Fisher Scientific) for 24 hours before storing in 70% ethanol. For µCT measurements at USDA (North Dakota), bones were scanned using a Scanco µCT scanner (µCT-40; Scanco Medical AG, Bassersdorf, Switzerland); together with reconstruction and 3D quantitative analyses, these experiments were performed as described previously (24).

### Primary osteoblast/osteoclast differentiation and coculture

Osteoblasts and osteoclasts were differentiated from mouse bone marrow cells as described previously (14). In brief, for osteoblast differentiation, bone marrow cells were obtained from adult *Erfe*^fl/fl^ and *Col2.3*-Cre;*Erfe*^fl/fl^ mice by flushing the bone marrow and bone marrow cells were cultured in αMEM (#15-012-CV, Corning) supplemented with 10% fetal bovine serum (#26140-079, Gibco) and 1% of antibiotic-antimycotic (#15240-062, Gibco) under 37°C in a humidified incubator with 5% CO_2_. Osteoblasts were then differentiated in the complete medium supplemented with 10 mM β-glycerophosphate (#35675, EMD Millipore), 50 µg/mL L-ascorbic acid (#AX1775-3, Supelco), and 10 nM dexamethasone (#D2916, Sigma). For osteoclast differentiation, bone marrow cells were cultured in αMEM supplemented with 50 ng/mL M-CSF (#315-02, PeproTech) for 48-72 hours. Osteoclasts were differentiated in the complete medium supplemented with 50 ng/mL M-CSF and 100 ng/mL RANKL (#315-11, PeproTech) for 7 days. Primary bone marrow cells from *Erfe*^+/+^ and *Erfe*^−/−^ mice were used for osteoblast:osteoclast co-culture experiments. Cells were seeded in transwell plate wells (#3450, Corning) and differentiated into osteoblasts as mentioned above. In parallel, osteoclasts were differentiated on transwell inserts. Once the cells were completed differentiation, the transwell inserts were placed into the wells and medium was changed to osteoblastic differentiation medium supplemented with 50 ng/mL M-CSF and 100 ng/mL RANKL. The plates were maintained at 37°C in a humidified incubator with 5% CO_2_ for 72 hours prior to analysis.

### Osteoblast mineralization and osteoclast differentiation measurement

Matrix mineralization formed by osteoblasts was determined by staining with 0.1% alizarin red (#A5533, Sigma) in 0.5% KOH. Tartrate-resistant acid phosphatase (TRAP) activity of osteoclasts was measured using TRAP staining kit (#387A, Sigma) according to the manufacturer’s instructions.

### RNA extraction and real-time PCR

Total RNA was extracted using RNAeasy Mini kit (#74106, Qiagen, Hilden, Germany) according to the manufacturer’s instructions. Complementary DNA was synthesized using a 500 ng of total RNA with High-Capacity cDNA Reverse Transcriptase kit (#4368814, ThermoFisher Scientific). Relative mRNA levels of genes of interest were determined using Power SYBER Green PCR Master Mix (#4367659, ThermoFisher Scientific) and measured on the QuantStudio 7 Pro Real-Time PCR (ThermoFisher Scientific). Primers are listed in Table S2.

### Statistics

All data are presented as means ± SEM. Statistical significance was assessed by two tail Student’s *t* test and a *P*-value of less than 0.05 (*P* < 0.05) was considered statistically significant.

## RESULTS

### *Erfe* Loss in Osteoblasts Does Not Affect Bone Formation

Given our prior findings demonstrating decreased bone mineral density (BMD) in *Erfe*^−/−^ mice (14), we initially compared bone mineral density (BMD) between *Erfe*^+/+^ (wild type), *Erfe*^−/−^, *Erfe*^fl/fl^ and *Col2.3*-Cre;*Erfe*^fl/fl^ mice using DXA (dual x-ray absorptiometry) scanning. We again confirm a decrease in BMD in *Erfe*^−/−^ mice when compared to *Erfe*^+/+^ but no difference in BMD between *Erfe*^fl/fl^ and *Col2.3*-Cre;*Erfe*^fl/fl^ mice in the whole body, femur, tibia, and lumbar L4-L6 vertebrae (Figure 1A). *Col2.3*-Cre;*Erfe*^*fl/fl*^ mice did not exhibit significant differences in the bone volume fraction (BV/TV) and trabecular thickness (Tb.Th) measurements based on calcium deposits in Von Kossa staining (Figures S1A and S1B). In addition, no significant differences were observed in bone formation characterized by mineral apposition rate (MAR) in the femur or vertebrae (Figures S1C and S1D). Furthermore, we observed no differences in BMD between *Col2.3*-Cre;*Erfe*^fl/fl^ and *Erfe*^fl/fl^ mice on the basis of gender (Figure S2A) or age (6 weeks old vs 14 weeks old) (Figure S2B). Finally, we confirmed by micro-CT analysis that *Erfe* loss in osteoblasts had no effect on BMD, BV/TV, Tb.Th, trabecular number (Tb.N), and trabecular spacing (Tb.Sp) when compared to *Erfe*^fl/fl^ mice (Figures 1B-1F). These data demonstrate that the functional role of ERFE in osteoblasts is dispensable for bone homeostasis at baseline.

**Figure 1.**
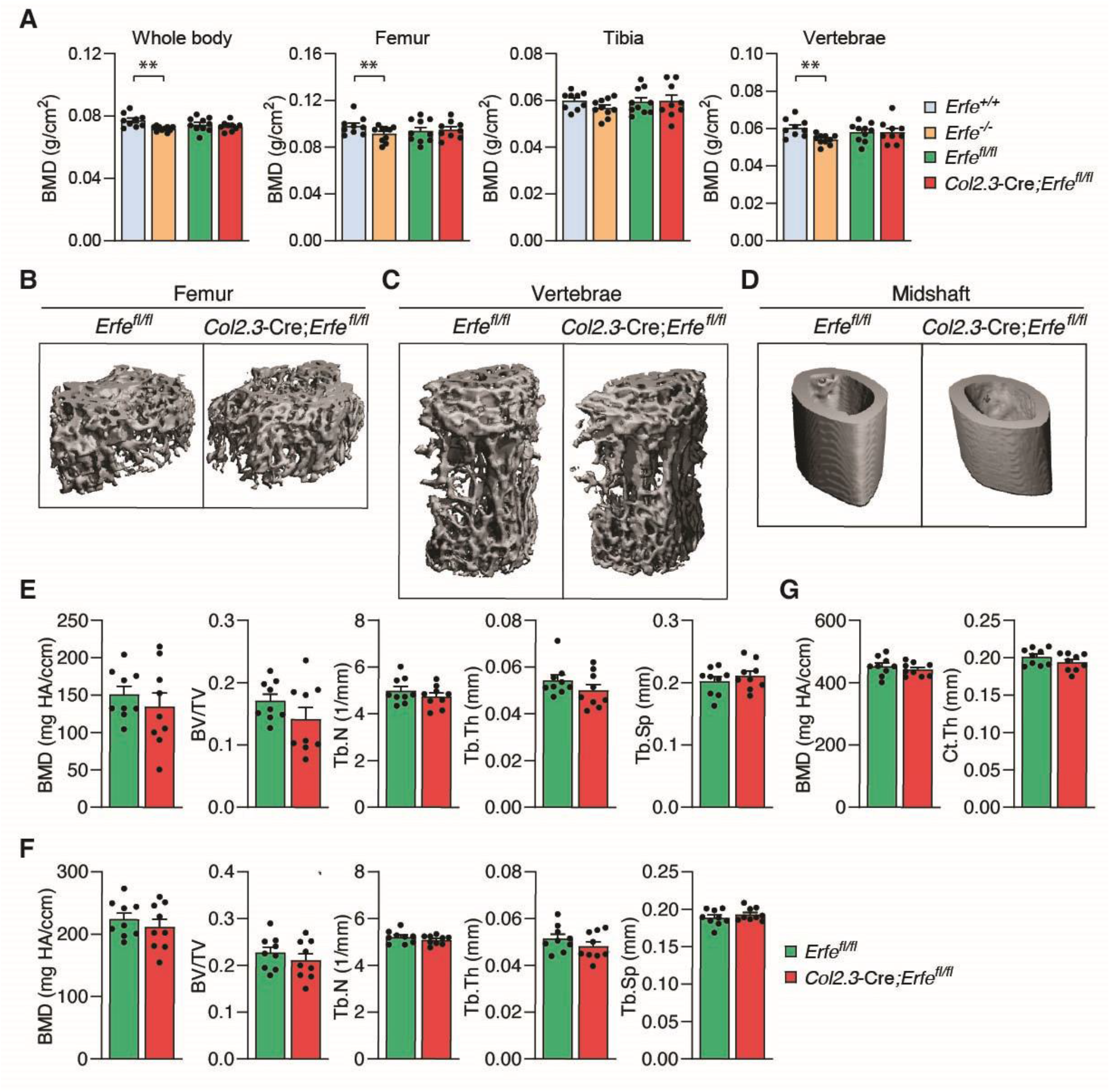
Erfe Loss in Osteoblasts Does Not Affect Bone Formation in Mice. (**A**) Bone mineral density (BMD) assessed using DXA scan in the whole body, femur, tibia, and lumbar vertebrae (L4-L6) of 14-week-old *Erfe*^+/+^ (wild type), *Erfe*^−/−^, *Erfe*^fl/fl^ and *Col2.3*-Cre;*Erfe*^fl/fl^ mice (*N* = 10). (**B-D**) Three-dimensional images of micro-CT analyses in the distal femur, lumbar vertebrae, and femoral midshaft. (**E-F**) Quantitative micro-CT analyses of trabecular structure; BMD, bone volume fraction (BV/TV), trabecular number (Tb.N), trabecular thickness (Tb.Th), trabecular separation (Tb.Sp) at the distal femur and lumbar vertebrae (*N* = 9). (**G**) Quantitative micro-CT analyses of cortical structure; BMD and cortical thickness (Ct.Th) at the midshaft of femoral bone (*N* = 9). Data are presented as mean ± SEM. \*\**P* < 0.01 using two-tail Student *t* test.

### *Erfe* Loss Accelerates Osteoblast Function *In Vitro*

We previously demonstrate that osteoblasts from *Erfe*^−/−^ mice exhibit enhanced mineralization and up-regulated biomarkers of osteoblast activity *in vitro* (14). Consistently, we confirm that osteoblasts from *Col2.3*-Cre;*Erfe*^fl/fl^ mice similarly exhibit increased mineralization as evidenced by alizarin red staining *in vivo* (Figure 2A). In addition, *in vitro* assessment of *Col2.3*-Cre;*Erfe*^fl/fl^ relative to *Erfe*^fl/fl^ osteoblasts demonstrates up-regulated biomarkers of osteoblast activity such as *Col1a1* and *Alp* (Figure 2B). In contrast, gene expressions of osteoblast and osteoclast biomarkers (*Runx2, Osx, Col1a1, Alp, Rankl, Opg, Acp5*, and *Ctsk*), *Bmp2* and *Bmp4* were unaffected in bone from *Col2.3*-Cre;*Erfe*^fl/fl^ relative to *Erfe*^fl/fl^ mice (Figure S3A and S3B). Finally, there were no differences of TRAP activity (Figure 2C) or osteoclast biomarker genes, *Acp5* and *Ctsk* (Figure 2D) in *Col2.3*-Cre;*Erfe*^fl/fl^ osteoclasts compared with *Erfe*^fl/fl^ osteoclasts *in vitro*.

**Figure 2.**
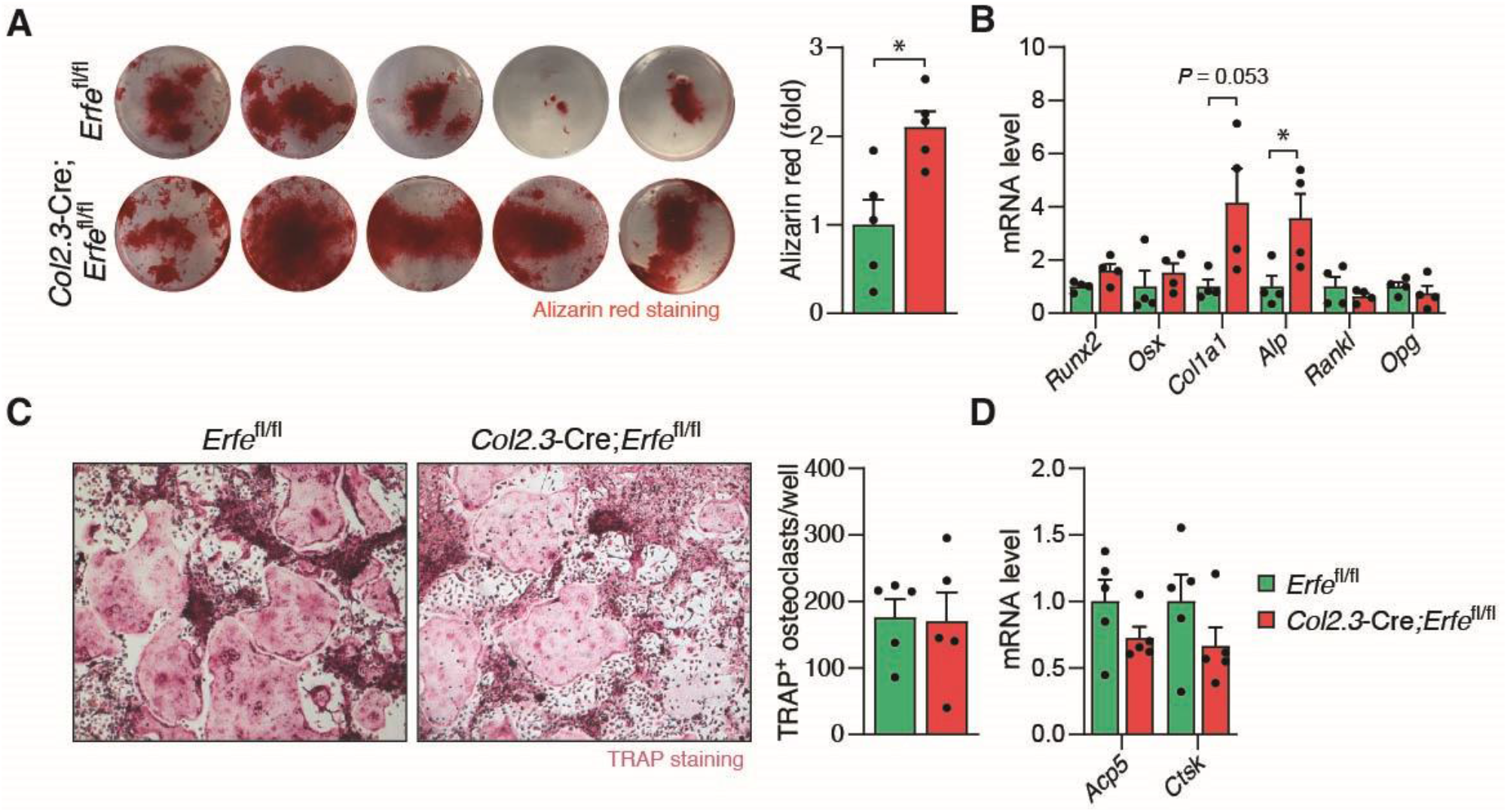
ERFE Loss Induces Osteoblast Activity. (**A**) Mineralization measuring by alizarin red staining of osteoblasts differentiated from the bone marrow of *Erfe*^fl/fl^ and *Col2.3*-Cre;*Erfe*^fl/fl^ mice (*N* = 5). (**B**) *Runx2, Osx, Col1a1, Alp, Rankl*, and *Opg* mRNA expression in *Erfe*^fl/fl^ and *Col2.3*-Cre;*Erfe*^fl/fl^ osteoblast culture (*N* = 4). (**C**) Representative images of TRAP staining for osteoclasts differentiated from the bone marrow of *Erfe*^fl/fl^ and *Col2.3*-Cre;*Erfe*^fl/fl^ mice (*N* = 5). (**D**) *Acp5* and *Ctsk* mRNA expression in *Erfe*^fl/fl^ and *Col2.3*-Cre;*Erfe*^fl/fl^ osteoclast culture (*N* = 5). Data are presented as mean ± SEM. \**P* < 0.05 using two-tail Student *t* test.

### Erfe loss in Osteoblasts Increases Bone Formation in Parathyroid Hormone Treated Mice

To assess whether enhanced activity in *Col2.3*-Cre;*Erfe*^fl/fl^ osteoblasts *in vitro* could affect dynamic skeletal homeostasis, we examined BMD in *Col2.3*-Cre;*Erfe*^fl/fl^ mice stimulated with parathyroid hormone (PTH) to induce bone formation and in ovariectomized (OVX) mice to induce bone resorption. Micro-CT scanning analysis demonstrates increased BMD in the trabecular bone, especially in vertebrae, but not in the cortical bone of PTH-treated *Col2.3*-Cre;*Erfe*^fl/fl^ relative to *Erfe*^fl/fl^ mice (Figure 3A-3F). A significant increase in BV/TV is detected in the trabecular bone in femur and vertebrae of PTH-treated relative to vehicle injected *Col2.3*-Cre;*Erfe*^fl/fl^ mice (Figure 3D and 3E). A significant increase in Tb.N is detected in the trabecular bone in femur but not in vertebrae of PTH-treated relative to vehicle injected *Col2.3*-Cre;*Erfe*^*fl/fl*^ mice; only a trend in increased Tb.N is detected in PTH-treated relative to vehicle in injected *Erfe*^fl/fl^ mice (Figures 3E). No altering in Tb.Th and Tb.Sp were observed (Figures 3F). Although PTH in general promotes preosteoblastic proliferation and osteoblastic bone formation (25-27), mechanistically, PTH acts as an anabolic agent by upregulating BMP signaling, particularly BMP2 and BMP4, to enhance mesenchymal stem cell differentiation into osteoblasts (28-31). We therefore anticipate that ERFE loss decreases BMP sequestration and increases BMP binding to receptors, leading to enhance osteoblast function (14). In contrast, following OVX, no significant differences in BMD between *Col2.3*-Cre;*Erfe*^fl/fl^ and *Erfe*^fl/fl^ mice were observed (Figures S4A-E). These findings indicate that *Erfe* loss in osteoblasts, possibly because of increased osteoblast activity, contributes to bone formation during anabolic but not catabolic stress.

**Figure 3.**
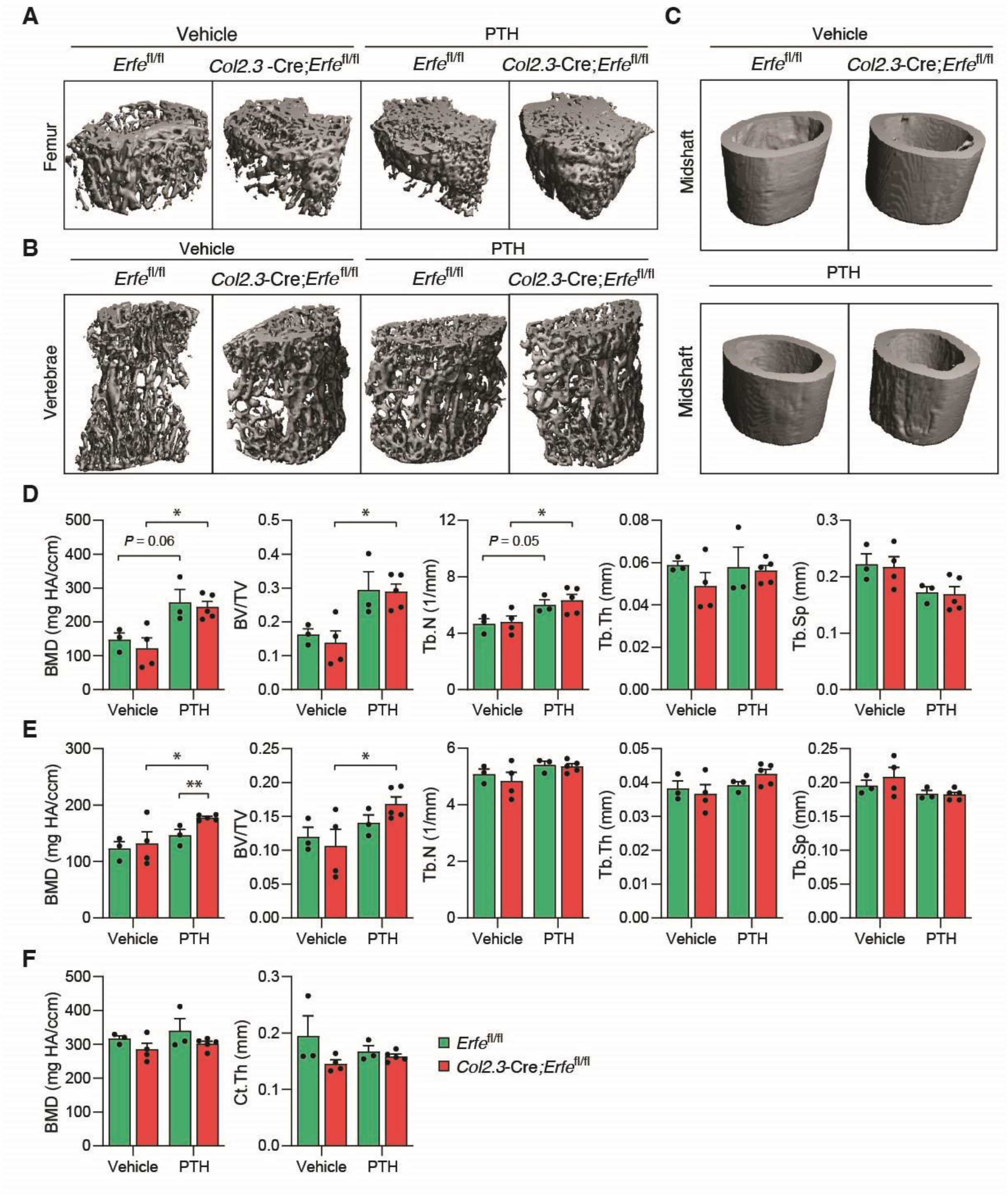
Osteoblastic *Erfe* Loss Promotes Bone Formation after Parathyroid Hormone Treatment in Mice. (**A-C**) Three-dimensional micro-CT images of the distal femur, lumbar vertebrae, and femoral midshaft from seven-week-old *Erfe*^fl/fl^ and *Col2.3*-Cre;*Erfe*^fl/fl^ mice (*N* = 3-5) treated with vehicle (0.9% NaCl) and 50 µg/kg parathyroid (PTH). (**D-E**) Quantitative micro-CT analyses of trabecular structure; BMD, bone volume fraction(BV/TV), trabecular number (Tb.N), trabecular thickness (Tb.Th), trabecular separation (Tb.Sp) at the distal femur and lumbar vertebrae from *Erfe*^fl/fl^ and *Col2.3*-Cre;*Erfe*^fl/fl^ mice after pTH treatment (*N* = 9). (**F**) Quantitative micro-CT analyses of cortical structure; BMD and cortical thickness (Ct.Th) at the midshaft of femoral bone of *Erfe*^fl/fl^ and *Col2.3*-Cre;*Erfe*^fl/fl^ mice after pTH treatment (*N* = 9). Data are presented as mean ± SEM. \**P* < 0.05, \*\**P* < 0.01 using two-tail Student *t* test.

### Osteoblastic ERFE Regulates Osteoblast:Osteoclast Crosstalk

Based on the reduction of BMD in *Erfe*^−/−^ mice, unchanged BMD in *Col2.3*-Cre;*Erfe*^fl/fl^ mice, and similarly increased activity in both *Erfe*^−/−^ and *Col2.3*-Cre;*Erfe*^fl/fl^ osteoblast *in vitro*, we hypothesize that the *in vivo* function of ERFE on bone involves osteoblast:osteoclast crosstalk. To test this hypothesis, we established osteoblast:osteoclast co-culture experiments (Figure S5A). These experiments demonstrate that expression of osteoblast activity biomarker genes *Runx2, Osx, Col1a1*, and *Alp* is increased in a similar manner in *Erfe*^−/−^ osteoblasts in non-co-cultured, co-cultured with *Erfe*^+/+^ osteoclasts, or co-cultured with *Erfe*^−/−^ osteoclasts conditions (Figure 4A), suggesting that increased osteoblast activity due to *Erfe* loss is only minimally affected by osteoblast:osteoclast crosstalk.

**Figure 4.**
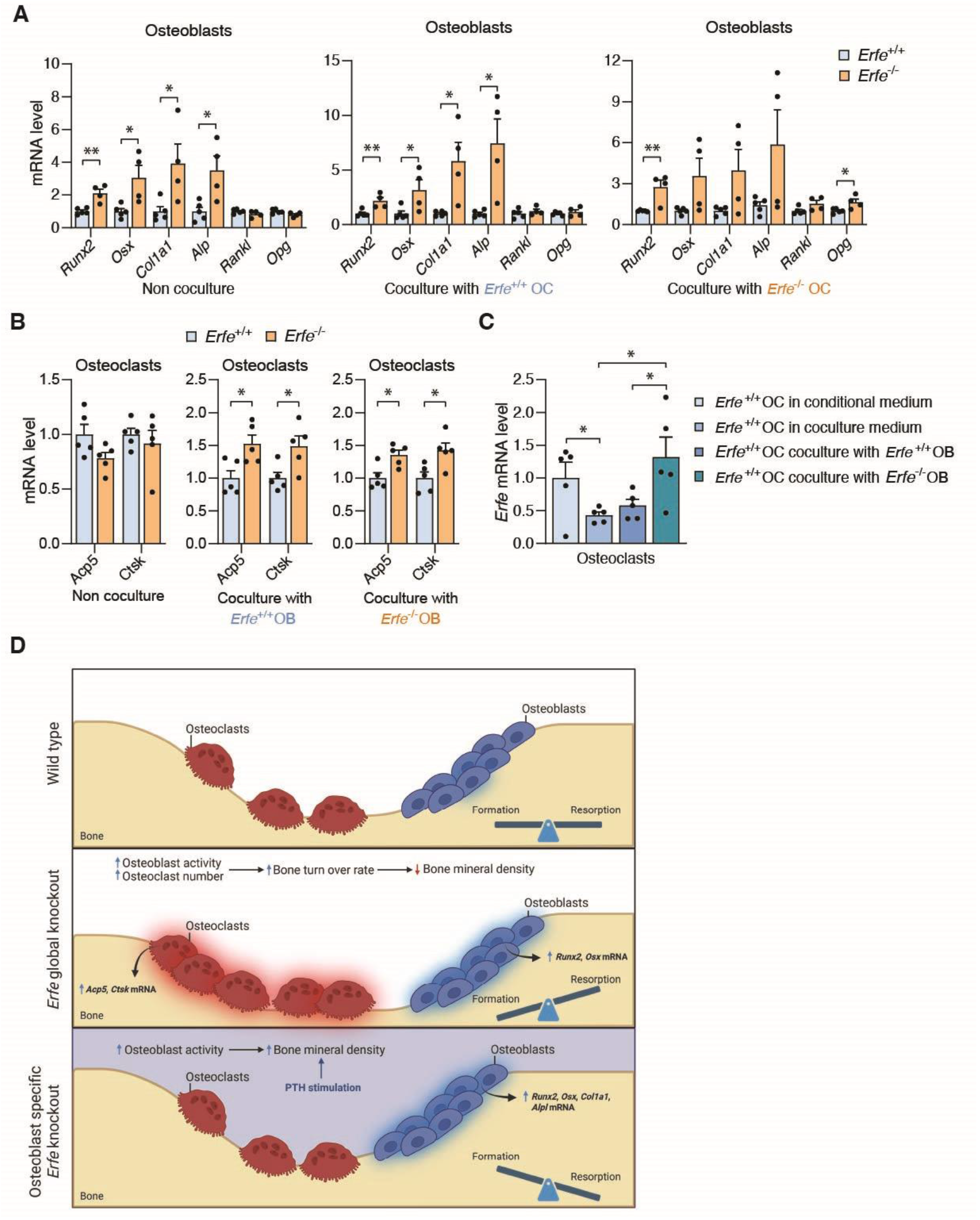
ERFE in Osteoclasts Regulates Osteoblast-Osteoclast Crosstalk. (**A**) mRNA levels of *Runx2, Osx, Col1a1, Alp, Rankl*, and *Opg* genes in *Erfe*^+/+^ and *Erfe*^−/−^ osteoblasts; non co-cultured, in co-culture with *Erfe*^*+/+*^ osteoclasts (OC), and in co-culture with *Erfe*^−/−^ osteoclasts (*N* = 4-5). (**B**) mRNA levels of *Acp5* and *Ctsk* genes in *Erfe*^+/+^ and *Erfe*^−/−^ osteoclasts; non co-cultured, in co-culture with *Erfe*^+/+^ osteoblasts (OB), and in co-culture with *Erfe*^−/−^ osteoblasts (*N* = 5). (**C**) *Erfe* expression in osteoclast (OC) cultured in condition medium, co-culture medium, co-cultured with *Erfe*^+/+^ osteoblasts (OB) and co-cultured with *Erfe*^−/−^ osteoblasts for 72 hours (*N* = 5). (**D**) Schematic presentation of the role of ERFE in osteoblasts and osteoclasts on the balance of bone formation and resorption. In normal condition, osteoblasts and osteoclasts work together to maintain healthy bone (Top). In contrast, lack of ERFE causes increased osteoblast and osteoclast activity leading to high rates of bone turnover, resulting in bone loss (Middle). In osteoblast-selective *Erfe* deficiency, osteoblasts (but not osteoclasts) exhibit increased activity, preventing bone loss (Bottom). Data are presented as mean ± SEM. \**P* < 0.05, \*\**P* < 0.01 using two-tail Student *t* test.

On the other hand, although there are no differences in expression of osteoclast activity genes *Acp5* and *Ctsk* between non-co-cultured *Erfe*^−/−^ or *Erfe*^+/+^ osteoclasts, *Acp5* and *Ctsk* expression is equivalently increased in *Erfe*^*−/−*^ osteoclasts co-cultured with *Erfe*^−/−^ or *Erfe*^+/+^ osteoblasts (Figure 4B). Normalized to non-co-cultured expression levels, *Erfe*^+/+^ osteoclasts showed decreased *Acp5* and *Ctsk* expression when co-cultured with *Erfe*^+/+^ and *Erfe*^−/−^ osteoblasts (Figure S5B), while *Erfe*^−/−^ osteoclasts showed increased *Acp5* and unchanged *Ctsk* expression (Figure S5C). In addition, expression of *Erfe* in *Erfe*^+/+^ osteoclasts is increased when co-cultured with *Erfe*^*−/−*^ osteoblasts (Figure 4C). Finally, although no differences are observed in BMP receptor gene *Alk3* in both co-cultured and non co-cultured osteoclasts, a significant increase is evident in *Bmpr2* only in co-cultured but not in non co-cultured cells (Figure S5D and S5E). Taken together, ERFE loss in osteoblasts enhances cell autonomous activity while ERFE loss in osteoclasts modulates osteoblast:osteoclast crosstalk to enhance osteoclast activity, providing supporting evidence for how selective ERFE loss in osteoblasts in *Col2.3*-Cre;*Erfe*^fl/fl^ mice does not enhance osteoclast activity and prevents the decreased BMD observed in *Erfe*^−/−^ mice (Figure 4D).

## DISCUSSION

ERFE normally functions as the erythroblast–derived regulator of iron metabolism that acts through hepcidin suppression during stress erythropoiesis (1). Our previous study uncovered a new role of ERFE in bone metabolism (14). Specifically, we showed that ERFE regulates BMP signaling in osteoblasts, limiting osteoclastogenesis with consequent decreased BMD in *Erfe*^−/−^ mice (14). More recently, we generated and characterized osteoblast-specific *Erfe* knockout mice (*Col2.3*-Cre;*Erfe*^fl/fl^), demonstrating that ERFE derived from bone–forming osteoblasts contributes to hepcidin regulation and red blood cell recovery during stress erythropoiesis (23). We now find that osteoblast-derived ERFE does not alone regulate BMD as *Col2.3*-Cre;*Erfe*^fl/fl^ mice do not recapitulate decreased BMD observed in *Erfe*^−/−^ mice, adding a new dimension to current knowledge. This differential effect on BMD may be dependent on normal (not increased) osteoclastogenesis in *Col2.3*-Cre;*Erfe*^fl/fl^ mice and enhanced *Erfe* expression in osteoclasts in *in vitro* co-culture experiments with osteoblasts. Of note is that the loss of osteoblast–derived ERFE in *Col2.3*-Cre;*Erfe*^fl/fl^ mice enhances BMD in response to PTH—positioning ERFE as a vital link between the osteoblast and the osteoclast during bone remodeling.

It is known that, during bone remodeling, multiple local cytokine circuits and systemic hormones regulate the delicate balance between bone resorption and bone formation. In fact, the paracrine and endocrine function of bone is well established, especially the evidence of communication between osteoblasts and osteoclasts. Osteoblasts build new bone and regulate osteoclastogenesis via production of RANKL (32-35). In addition, OPG, WNT inhibitor secreted Frizzled-related protein 1, and LGR4 all provide mechanisms for titrating RANKL to alter the net balance between bone formation and bone resorptions (36-40). Finally, bone remodeling is regulated by BMPs, BMP receptors, and Noggin (NOG) expressed in osteoblasts. Noggin is secreted by osteoblasts, binds to and competitively blocks the action of BMPs (i.e. BMP2 and BMP4). Evidence from osteoblast-selective *Nog* overexpressing mice, with reduced bone formation and osteopenia, support an important role for the BMP pathway in adult bone forming capacity (41). While osteoinduction by BMPs has long been used to accelerate fracture healing, using BMP-related therapeutics to increase BMD is complicated by ectopic ossification. Our current data provides an additional opportunity to dissect how ERFE-mediated modulation of BMPs may direct bone remodeling.

Global loss of ERFE results in decreased BMD (14). In contrast, we find that the deletion of ERFE selectively in osteoblasts in *Col2.3*-Cre;*Erfe*^fl/fl^ mice does not decrease BMD and enhances BMD after PTH. This suggests that osteoblast–derived ERFE acts to restrict bone formation. This novel regulatory mechanism for bone homeostasis is consistent with *Erfe* expression *in vitro* in osteoblasts and *ex vivo* in whole bone, evidence for functional ERFE in osteoblast–conditioned media, and enhanced mineralization of *Erfe*^−/−^ osteoblasts *in vitro* (14). Mechanistically, we show in additional *in vitro* osteoblast:osteoclast co-culture experiments that the osteoblastogenesis gene program is enhanced in response to ERFE loss in osteoblasts and is unaffected by control or *Erfe*^−/−^ osteoclasts. Previous studies of osteoblast:osteoclast crosstalk in co-culture showing that TRAP activity in osteoclasts is reduced by the presence of osteoblasts (42). However, the osteoclastogenesis gene program is enhanced in response to ERFE loss in osteoclasts in co-culture experiments and is not further affected by ERFE loss in osteoblasts. Taken together, our findings unmask a new mechanism for osteoclastogenesis regulation that is dependent on osteoclast–derived ERFE as an important regulator of osteoblast:osteoclast crosstalk and confirms that decreased BMD in response to global ERFE loss is a consequence specifically of osteoclast ERFE loss which supports enhanced osteoclastogenesis in crosstalk between osteoblasts and osteoclasts.

## ACKNOWLEDGEMENTS

We sincerely appreciate Tomas Ganz and Elizabeta Nemeth (University of California, Los Angeles), as well as Martina Rauner (University of Dresden) for many stimulating and helpful discussions. We thank the Genome Engineering and iPSC Center at Washington University in St. Louis for their gRNA and donor design and validation services in making the *Erfe*^fl/fl^ mouse. T.Y., M.Z., and Y.Z.G. acknowledge the support of the National Institute of Diabetes and Digestive and Kidney Diseases (NIDDK) (R01 DK107670). Y.Z.G. additionally acknowledges the support of NIDDK (R01 DK095112). The authors also acknowledge R01 AG071870 to M.Z., and T.Y.; R01 AG074092, U01 AG073148 and R61 AG094602 to T.Y. and M.Z.; and U19 AG060917 and R01 DK113627 to M.Z.

## AUTHORSHIP CONTRIBUTIONS

P.N. and G.V.C. designed research, performed research, analyzed data, and wrote and edited the paper; M.P., E.T, A.G., G.P., R.W., E.I., F.S., F.K., and M.L. performed research, analyzed data, and edited the paper; J.C., and L.K. contributed vital reagents or analytical tools and edited the paper; T.Y., M.Z., and Y.Z.G. designed research, contributed vital reagents or analytical tools, analyzed data, and wrote and edited the paper.

