## Supplementary information for "Erythroferrone Modulates Osteoblast-Osteoclast Crosstalk During Bone Remodeling"

Figures S1 to S5

Tables S1 and S2

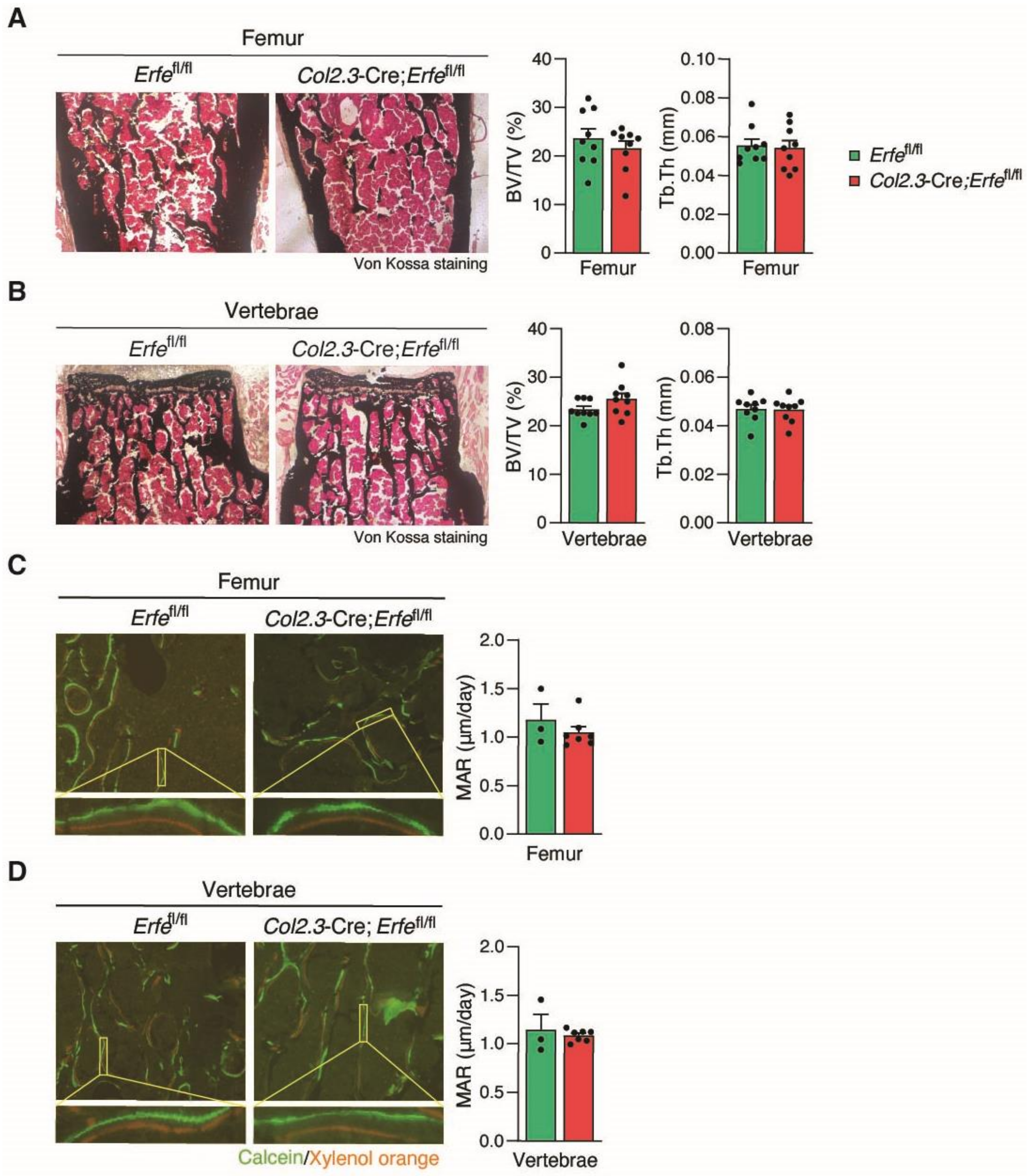


__Figure S1__: **Unchanged Calcium Deposits and Appositional Bone Growth in Osteoblast-Selective *Erfe* Knockout Mice.** (**A**-**B**) Von Kossa staining images of the femur and lumbar vertebrae and its histomorphometric measurements for bone volume fraction (BV/TV) and trabecular thickness (Tb.Th) from 14-week-old *Erfe*$^{fl/fl}$ and *Col2.3*-Cre;*Erfe*$^{fl/fl}$ male mice (*N* = 9). (**C**-**D**) Fluorescent images of mineral apposition rate (MAR) determined by the distance between the lines of mineral deposition from calcein green and xylenol orange injections (*N* = 3-5). Data are presented as mean ± SEM.

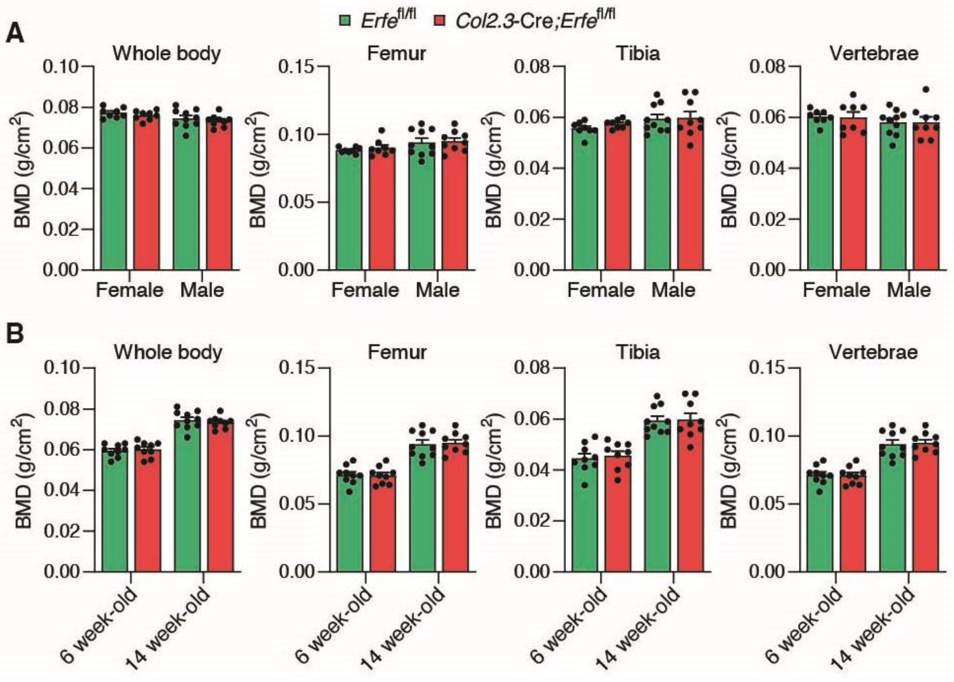


**Figure S2: Osteoblast-Selective *Erfe* Knockout Mice Do Not Exhibit Age- and Gender-Specific Differences in Bone Characteristics.** (**A**) Comparison of bone mineral density (BMD) in the whole body, femur, tibia, and lumbar vertebrae (L4-L6) measured by DXA scan between 14-week-old *Erfe*$^{fl/fl}$ and Col2.3-Cre;*Erfe*$^{fl/fl}$ male and female mice (*N* = 9). (**B**) Comparison of BMD between 6-week-old and 14-week-old *Erfe*$^{fl/fl}$ and Col2.3-Cre;*Erfe*$^{fl/fl}$ male mice (*N* = 9). Data are presented as mean ± SEM.

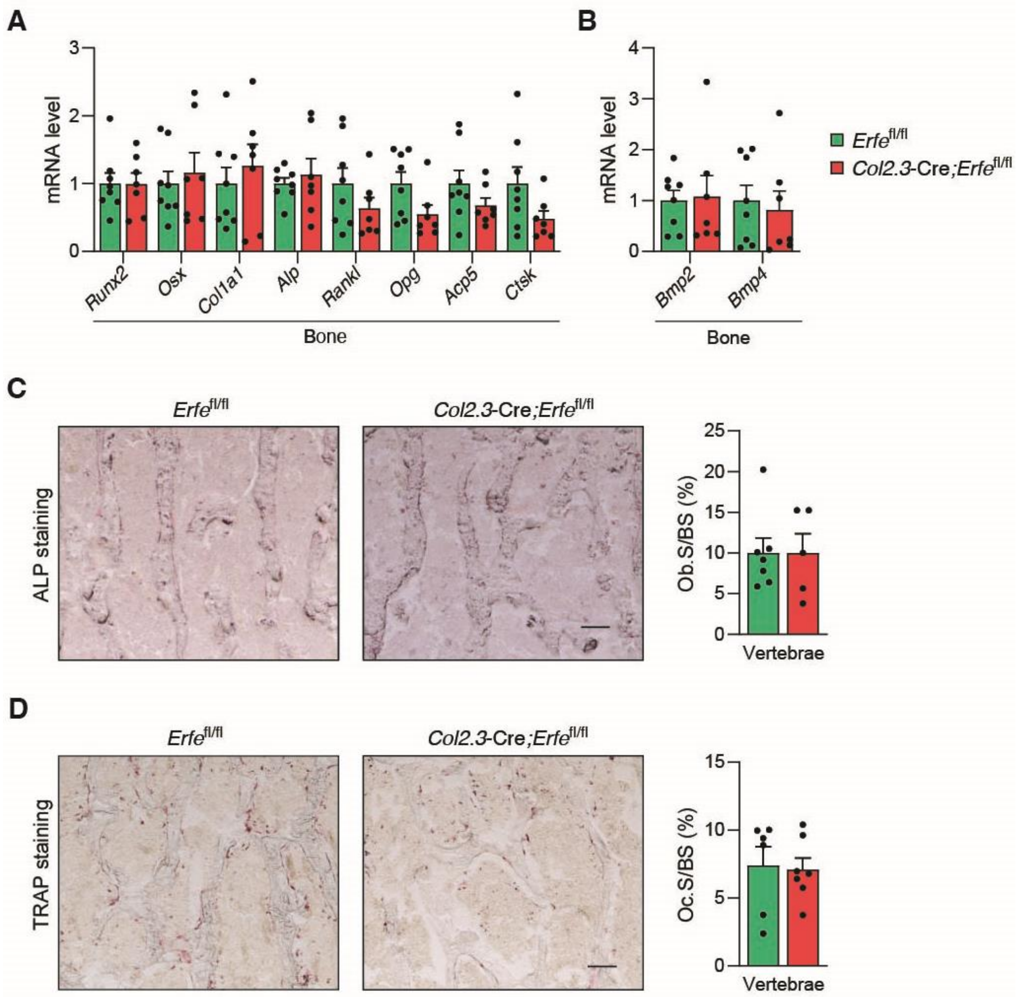


**Figure S3**: ***Erfe* Loss in Osteoblasts Does Not Affect Gene Expression and Cellularity of Osteoblasts and Osteoclasts in the Bone *In Vivo*.** (**A**) mRNA levels of osteoblast and osteoclast biomarker genes; *Runx2*, *Osx*, *Col1a1*, *Alp*, *Rankl*, *Opg*, *Acp5* and *Ctsk* in the bone of 14-week-old *Erfe*$^{fl/fl}$ and Col2.3-Cre;*Erfe*$^{fl/fl}$ mice (*N* = 3-4). (**B**) *Bmp2* and *Bmp4* mRNA levels in the bone of *Erfe*$^{fl/fl}$ and *Col2.3*-Cre;*Erfe*$^{fl/fl}$ mice (*N* = 3-4). (**C**-**D**) Representative images of ALP and TRAP staining and percentage of osteoblast and osteoclast surface determined by bone morphometric analysis of the vertebrae from 14-week-old *Erfe*$^{fl/fl}$ and *Col2.3*-Cre;*Erfe*$^{fl/fl}$ mice (*N* = 5-7). Scale bar 0.1 mm. Data are presented as mean ± SEM.

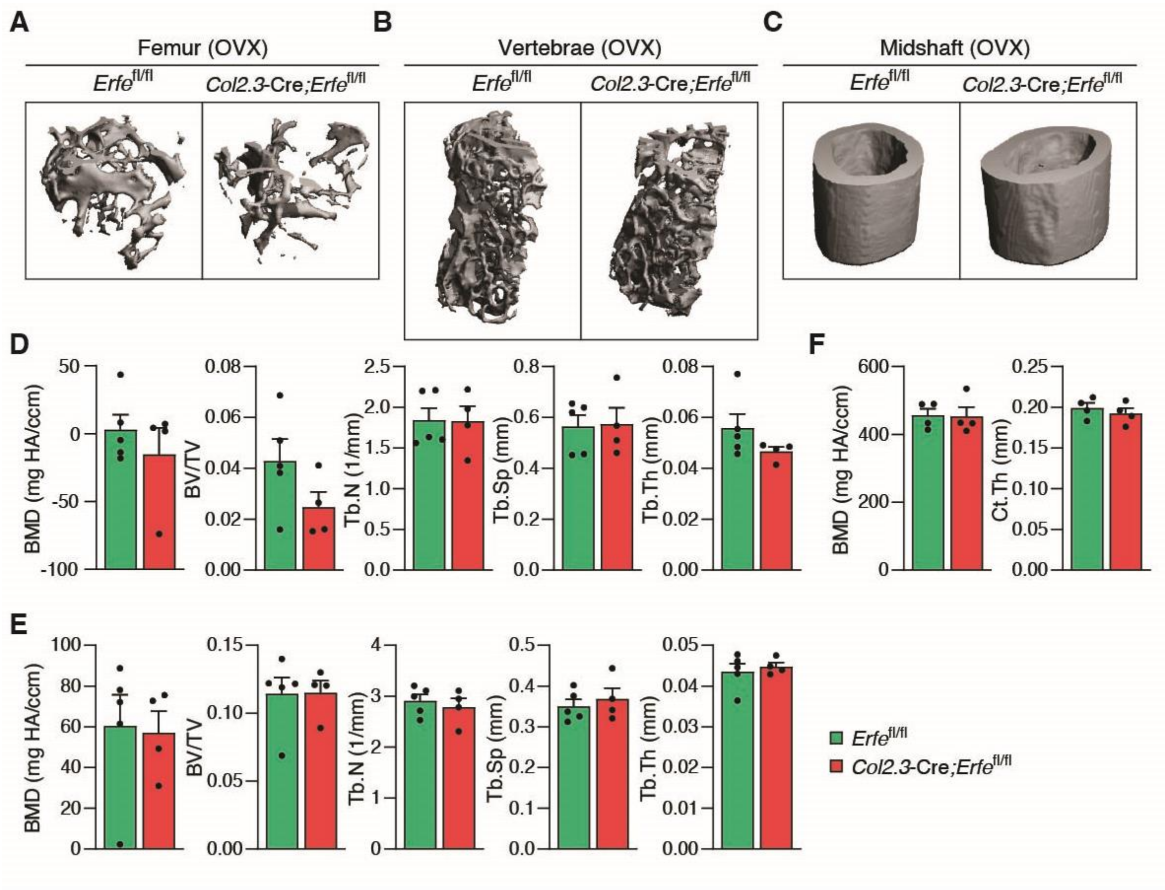


**Figure S4: Micro-Architecture Analysis of the Trabecular and Cortical Bone in Ovariectomized Osteoblast-Selective *Erfe* Knockout Mice.** (**A**-**C**) Three-dimensional micro-CT images of the femur, lumbar vertebrae, and femoral midshaft of 40-week-old *Erfe*$^{fl/fl}$ and *Col2.3*-Cre; *Erfe*$^{fl/fl}$ female mice following ovariectomy (OVX). (**D-E**) Bone mineral density (BMD), bone volume fraction (BV/TV), trabecular number (Tb.N), trabecular thickness (Tb.Th), and trabecular separation (Tb.Sp) of the femur and lumbar vertebrae from *Erfe*$^{fl/fl}$ and *Col2.3*-Cre; *Erfe*$^{fl/fl}$ mice (*N* = 4-5). (**F**) BMD and cortical thickness (Ct.Th) of femoral midshaft from *Erfe*$^{fl/fl}$ and *Col2.3*-Cre; *Erfe*$^{fl/fl}$ mice (*N* = 4). Data are presented as mean ± SEM.

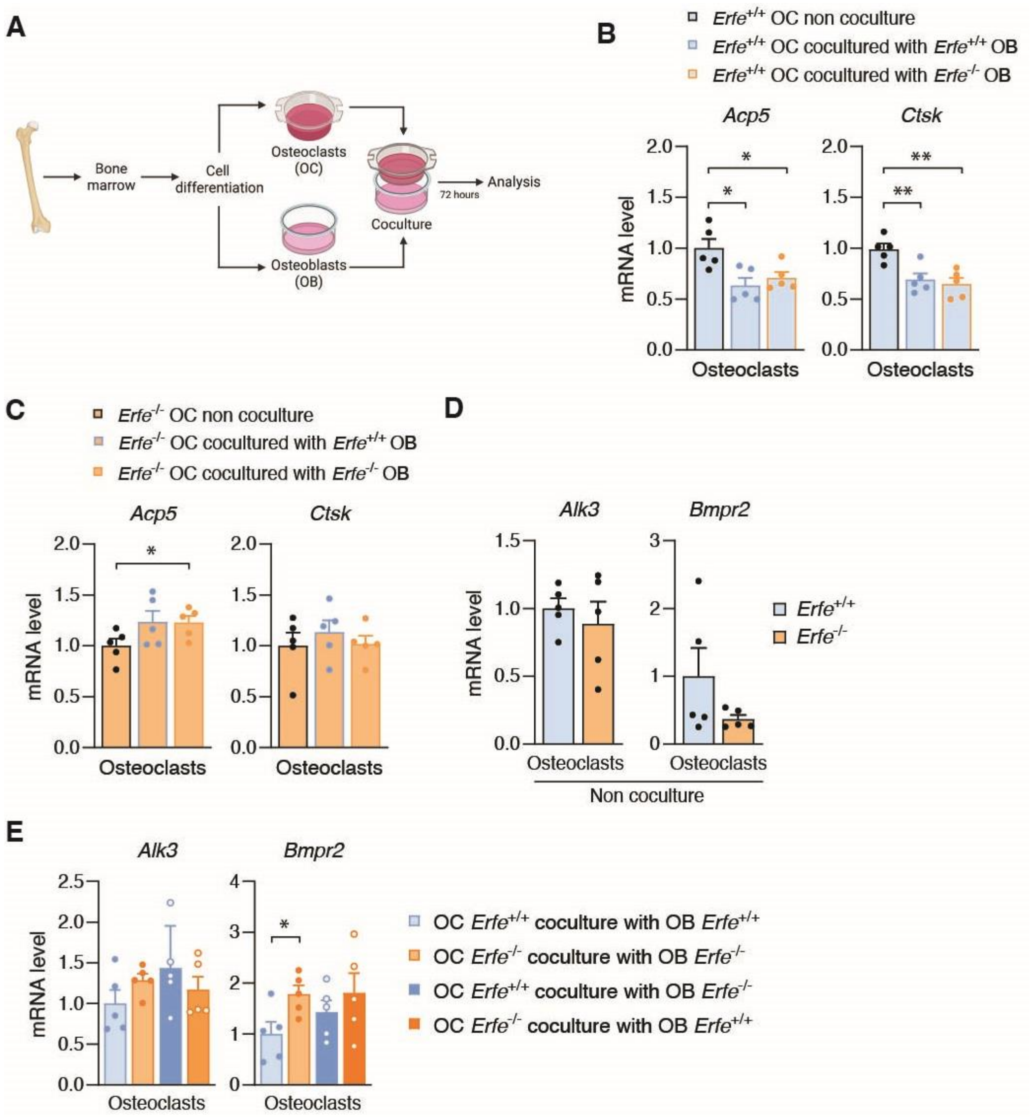


<u>Figure S5</u>: ***Erfe* Loss in Osteoblast Promotes Osteoclastogenesis Gene Expression During Osteoblast:Osteoclast Crosstalk in Co-culture.** (**A**) Schematic illustrating experimental setup for indirect osteoblast:-osteoclast co-cultures. Mature osteoblasts and osteoclasts were co-cultured with no cell-cell contact for 72 hours using transwell inserts. (**B-C**) *Acp5* and *Ctsk* expression in *Erfe*$^{+/+}$ osteoclasts and *Erfe*$^{-/-}$ osteoclasts under non co-cultured conditions, co-cultured with *Erfe*$^{+/+}$ osteoblasts (OB), and co-cultured with *Erfe*$^{-/-}$ OB ($N$ = 5). (**D**) mRNA levels of BMP receptor genes *Alk3* and *Bmpr2* in osteoclasts non coculture ($N$ = 5). (**E**) *Alk3* and *Bmpr2* expression of *Erfe*$^{+/+}$ and *Erfe*$^{-/-}$ osteoclasts co-cultured with *Erfe*$^{+/+}$ or *Erfe*$^{-/-}$ osteoblasts ($N$ = 5). Data are presented as mean ± SEM. *$P$ < 0.05, **$P$ < 0.01 using two-tail Student $t$ test.

### Tables

#### Table S1: PCR primers for genotyping

| Primer name | Sequence | Product size |
|---|---|---|
| *Primer set 1* *(the 5' loxP site)* | Forward:5'-TATGTGTGTGTGGTCAGCCC-3'<br>Reverse:5'-GGAGGTCCTTAGGGCTCTCA3' | 394 bp Floxed *Erfe*<br>354 bp WT |
| *Primer set 2* *(the 3' loxP site)* | Forward:5'-CCAGGCCCCTTTATCCCATC-3'<br>Reverse:5'-ACAGAAGCAGGAAAGGGCTC-3' | 400 bp Floxed *Erfe*<br>360 bp WT |
| Universal-cre | Forward:5'-CAAGTGACAGCAATGCTGTTTCAC-3'<br>Reverse:5'-CAGGTATCTCTGACCAGAGTCATC-3' | 550 bp |
| *Erfe*$^{-/-}$ | Forward:5'-GCAGCGCATCGCCTTCTATC-3'<br>Reverse:5'-GACCGTCACTGAGGTTCCAC-3' | 390 pb |
| *Erfe*$^{+/+}$ | Forward:5'-GTCAGCCTTACCTGCCCAG-3'<br>Reverse:5'- GACGTGAATCTCAGTCTGGC-3' | 216 pb |

#### Table S2: Oligonucleotide primers for Real-time qPCR

| Gene name | Forward (5'−3') | Reverse (5'−3') | Reference |
|---|---|---|---|
| *Erfe* | ATGGGGCTGGAGAACAGC | TGGCATTGTCCAAGAAGACA | (1) |
| *Acp5* | ACCTGTGCTTCCTCCAGGAT | TCTCAGGGTGGGAGTGGG | (2) |
| *Runx2* | GTGGCCACTTACCACAGAGC | GTTCTGAGGCGGGACACC | (2) |
| *Alp* | ACACCTTGACTGTGGTTACTGCTGA | CCTTGTAGCCAGGCCCGTTA | (2) |
| *Osx* | TGAGGAAGAAGCCCATTCAC | GTGGTCGCTTCTGGTAAAGC | (2) |
| *Col1a1* | CCTGGCAAAGACGGACTCAAC | GCTGAAGTCATAACCGCCACTG | (2) |
| *Rankl* | CAGCCATTTGCACACCTCAC | GTCTGTAGGTACGCTTCCCG | (2) |
| *Opg* | ACAGTTTGCCTGGGACCAAA | CAGGCTCTCCATCAAGGCAA | (2) |
| *Bmp2* | GCGCAGCTTCCATCACGAAG | ATTGAAGAAGAAGCGCCGGG | This study |
| *Bmp4* | TCCGTAGTGCCATTCGGAGC | GCCTCCTAGCAGGACTTGGC | (3) |
| *Bmpr1a* | CCCGATTTATGAAAATATGCATCGC | GCAATGACTTTCACCTGCTGCT | This study |
| *Bmpr2* | AGCTGCTGCTTCCTAGCTACTAC | ACACTTACCAGGTCTGTGGCTT | This study |
| *Hprt* | GCAGTCCCAGCGTCGTGATT | GCCACAATGTGATGGCCTCC | This study |
| *Actin* | TTCTTTGCAGCTCCTTCGTT | ATGGAGGGGAATACAGCCC | (2) |

### SI Materials and Methods

#### *Histomorphometry analysis*

Mice received intraperitoneal injection of 15 mg/kg calcein (#C0875, Sigma) at day -8 and 90 mg/kg xylenol (#398187, Sigma) at day -2 prior to sacrifice. Femur and lumbar vertebra were dissected and immediately fixed in 10% neutral buffered formalin. After 24 h, bones were washed under running water for 30 min and immersed in 30% sucrose (#S0389, Sigma) in PBS for 24-72 h at 4°C. Bones were then embedded in a cryomold with OCT (#23-730-571, Fisher Scientific), prior to section at -25°C. The distance between xylenol and calcein lines in the bone sections were measured to yield mineral apposition rate (MAR). Von Kossa staining kit (#CSC0125P, StatLab) was used to measure trabecular bone volume fraction (BV/TV, %) and trabecular thickness (Tb.Th) based on calcium deposits. Sections were visualized under the Zeiss AXIO observer Z1 inverted microscope. Histopathological images were analyzed using QuPath 0.5.
